# TREM2 drives accumulation of pro-scarring monocyte-derived macrophages in the infarcted myocardium

**DOI:** 10.64898/2026.08.14.744182

**Authors:** Giuseppe Rizzo, Marie Piollet, Tobias Krammer, Ecem T. Sakalli, Alexander M. Leipold, Julius Gropper, Paul Alayrac, Adrien Tin-Kin-Wang, Manuel Gendre, Thomas A. Prohaska, Anahi Paula Arias-Loza, Ludovica Timperi, Anna Rizakou, Sourish Reddy Bandi, Dirk J.J. Schulz, Andrea Ninni, Daniele Lettieri-Barbato, Marco Colonna, Christopher K. Glass, Jean-Sébastien Silvestre, Stephane Camus, Alma Zernecke, Antoine-Emmanuel Saliba, Clément Cochain

## Abstract

Myocardial infarction is a leading cause of death and disability worldwide. Ischemic injury leads to irreversible loss of cardiomyocytes, the contractile cells of the heart, and formation of a fibrotic scar. After infarction, macrophages massively infiltrate the heart and orchestrate the tissue repair process by removing dead cells and modulating fibroblast activation for scar formation. We previously demonstrated that diverse monocyte-derived macrophage populations dynamically accumulate in the heart following myocardial infarction, notably a pro-repair *Trem2^hi^* subset. In this study, we leveraged spatial transcriptomics, single-cell RNA-seq, and functional assays to elucidate the role of TREM2 in driving macrophage-mediated cardiac tissue repair post-infarction. We show that *Trem2^hi^* macrophages localize in scarring areas of the infarcted myocardium in the vicinity of collagen-producing myofibroblasts. In *Trem2^-/-^* mice, cardiac accumulation of monocyte-derived macrophages with a pro-scarring ‘matrisome-associated macrophage’ signature was reduced. TREM2 deficiency was functionally associated with reduced fibroblast proliferation, accumulation of myofibroblasts, decreased collagen deposition in the infarcted heart, and increased infarct size. *In vitro*, we show that TREM2 mediates efferocytosis-induced pro-fibrotic gene expression and promotes macrophage ability to induce fibroblast migration. IL-4 priming of bone marrow-derived macrophages further increased the pro-fibrotic response in macrophages, suggesting that IL-4 and efferocytosis act synergistically to drive this phenotype. Altogether, our results show that TREM2 is essential for the accumulation and function of pro-scarring monocyte-derived macrophages in the infarcted myocardium.

## Introduction

Macrophages have a critical and dual role in cardiac repair after myocardial infarction (MI). While these cells perform functions essential for tissue healing, such as clearing dead cells and debris from the ischemic wound and orchestrating scar formation to preserve myocardial structural integrity, they can concurrently exacerbate tissue damage, promote interstitial fibrosis, and ultimately impair cardiac function (1). The healthy and infarcted heart contain heterogeneous and ontogenically diverse macrophage populations, comprising self-renewing tissue resident macrophages, and macrophages differentiated from infiltrating monocytes (2). While tissue-resident macrophages are widely recognized as cardioprotective, monocyte-derived populations (frequently defined by CCR2 expression) are traditionally viewed as deleterious (2). However, recent evidence highlights that monocyte-derived macrophages comprise highly heterogeneous subpopulations with distinct, specialized functions (3); notably, certain subsets have been shown to exert protective roles, such as preventing post-infarction arrhythmias (4). Previously, we characterized the dynamics of monocyte-derived macrophage heterogeneity in experimental myocardial infarction, identifying macrophage subsets enriched for the expression of *Trem2*, that we termed *Trem2^hi^* macrophages(3). Cardiac *Trem2^hi^* monocyte-derived macrophages displayed a rather ‘alternatively activated’ macrophage profile with low expression of inflammatory cytokines and expression of genes involved in wound healing and scar formation, such as *Spp1* and *Gpnmb* (3). Macrophages with a similar gene expression signature were observed in the human ischemic heart (3), (5).

Myocardial *Trem2^hi^* monocyte-derived macrophages express a core signature including *Trem2*, *Cd9*, *Spp1*, *Gpnmb*, *Fabp5*, *Cd63* characteristic of a conserved, disease-associated macrophage subset identified across various tissues and clinical contexts. This phenotype closely mirrors Lipid-= Associated Macrophages (‘LAM’) in obesity (6), non-alcoholic steatohepatitis (NASH)/non-alcoholic fatty liver disease NAFLD (7), (8), (9), and atherosclerosis (10), Scar Associated Macrophage (‘SAM’) in the fibrotic liver (11), fibrogenic ‘Fab5’ macrophages in the lung and liver (12), and Disease-Associated Microglia (‘DAM’) in the central nervous system (13). Several reports associated the presence of such macrophages with tissue fibrosis in both mice and human, including in lung of COVID-19 patients with acute respiratory distress syndrome (14), in the liver with metabolic dysfunction-associated steatohepatitis (MASH) (15), or NASH (9). Cross-tissue analysis of this macrophage subset furthermore identified a Matrisome-Associated Macrophages (MAM) core signature in human fibrosis-associated diseases (16). Moving forward, we will refer to this population of macrophages as *Trem2^hi^*.

In several pathological contexts, the accumulation of macrophages exhibiting such *Trem2^hi^* signature has been shown to depend on Triggering Receptor Expressed on Myeloid cells-2 (TREM2) (17). TREM2 is a receptor expressed by mononuclear phagocytes that regulates key functions of macrophages including survival, cytokine production, lipid metabolism and efferocytosis (17). TREM2 controls macrophage function in various cardiometabolic diseases such as NAFLD (9), obesity (6) and atherosclerosis (18), (19). In the heart, TREM2 has been ascribed protective functions in experimental viral myocarditis (20) and experimental hypertensive heart failure (21). Recruited monocyte-derived TREM2^+^ macrophages have been shown to promote atrial fibrillation via *Spp1*-mediated tissue remodeling (22). In experimental MI, application of recombinant soluble TREM2 was linked to improved cardiac function (23). Additionally, TREM2-mediated efferocytosis induced metabolic rewiring of macrophages and production of itaconate, inhibiting cardiomyocytes apoptosis and promoting fibroblasts proliferation after MI (24).

In this study, we demonstrate that post-infarction myocardial *Trem2^hi^*macrophages are highly enriched in a pro-fibrotic, matrisome-associated gene signature and directly interface with collagen-producing myofibroblasts. Genetic deletion of *Trem2* impaired the accumulation of macrophages expressing this matrisome-associated program; consequently, *Trem2*^-/-^ mice exhibited reduced interstitial fibrosis but developed larger, maladaptive scars four weeks post-myocardial infarction. Mechanistically, *in vitro* assays revealed that efferocytosis drives the acquisition of this pro-fibrotic phenotype in a TREM2-dependent manner. Taken together, our findings identify TREM2 as a critical regulator of efferocytic, pro-scarring macrophage accumulation, orchestrating the balance between definitive scar formation and adverse interstitial fibrosis in the ischemic myocardium

## Methods

Additional detailed methods can be found in the **Supplementary Materials.**

### Mouse myocardial infarction

Myocardial infarction was induced by permanent ligation of the Left Anterior Descending (LAD) coronary artery in 8- to 12-week-old *Trem2^+/+^* and *Trem2^-/-^* male mice on a C57BL6/J background, under isoflurane anesthesia (5% induction; 2-2.5% maintenance) and buprenorphine analgesia (0.1mg/kg; s.c.). Analgesia was maintained in the two days following surgery with twice daily administration of Buprenorphine (0.1mg/kg, s.c.). All animal studies conform to the Directive 2010/63/EU of the European Parliament and have been approved by the appropriate local authorities (Regierung von Unterfranken, Wuerzburg, Germany, Akt.-Z. 55.2-DMS-2532-2-743, Akt.-Z. 55.2.2-2532.2-865, Akt.-Z. 55.2.2-2532-2-1594).

### Spatial and scRNA-seq analyses

Spatial transcriptomics of an infarcted mouse heart (day 7 post-MI) was performed on a fresh frozen cryosection using the Curio Seeker Spatial Mapping Kit (Curio Biosciences), alongside histopathological staining of the consecutive section. ScRNA-seq libraries were generated using the 10x Genomics Chromium Next GEM Single Cell 3’kit v3. Data were analyzed in R v4.5.1 using the R packages Seurat v5.3.0 (25), SeuratExtend 1.2.5 (26), spacexr v2.2.1 (27), and ClusterFoldSimilarity v1.7.1 (28). All the single-cell and spatial transcriptomics data newly generated for this report will be made available upon publication.

### Statistical analysis

Statistical analyses were performed using GraphPad Prism version 10. Results are expressed as mean ± s.e.m. For two-group comparisons, normal distribution of the data was assessed by a D’Agostino–Pearson test followed by an unpaired t-test (normally distributed data) or a non-parametric Mann–Whitney test (non-normally distributed data). Data with multiple comparisons were assessed by one-way ANOVA followed by a Holm-Šídák’s multiple comparisons test. P values less than 0.05 were considered statistically significant.

## Results

### *Trem2^hi^* macrophages co-localize with myofibroblasts in the infarcted heart

To map the post-MI cardiac cellular landscape, we performed single-cell RNA sequencing (scRNA-seq) on murine hearts at baseline (uninjured) and at days 5, 7, and 14 post-MI, alongside non-immune cellular populations at baseline and day 7 post-MI (**Figure 1a and Supplemental Figure 1A**). MI induced a marked shift in cardiac cellular composition, characterized by recruitment of immune cells, such as monocyte-derived macrophages, neutrophils, dendritic cells, T and NK cells, as well as expansion and activation of fibroblasts (**Figure 1b-c**), consistent with previous studies (3), (29). Macrophages are widely recognized as critical players in fibrosis across multiple organs and diseases (16), (30). Analysis of cardiac macrophages revealed three main (‘Macro Resident’, ‘Macro *Trem2*’ and ‘Macro MHCII’) and three minor clusters (Macro ‘*Pde4c*’, ‘Macro *Isg15*’ and ‘Macro *Fn1*/*Ltc4’*”) in the infarcted heart (**Figure 1d**). *Trem2* expression was restricted to macrophages and particularly enriched in a macrophage population here termed ‘Macro *Trem2*’ (**Figure 1d**, **Supplementary Figure 1A)**.

**Figure 1.**
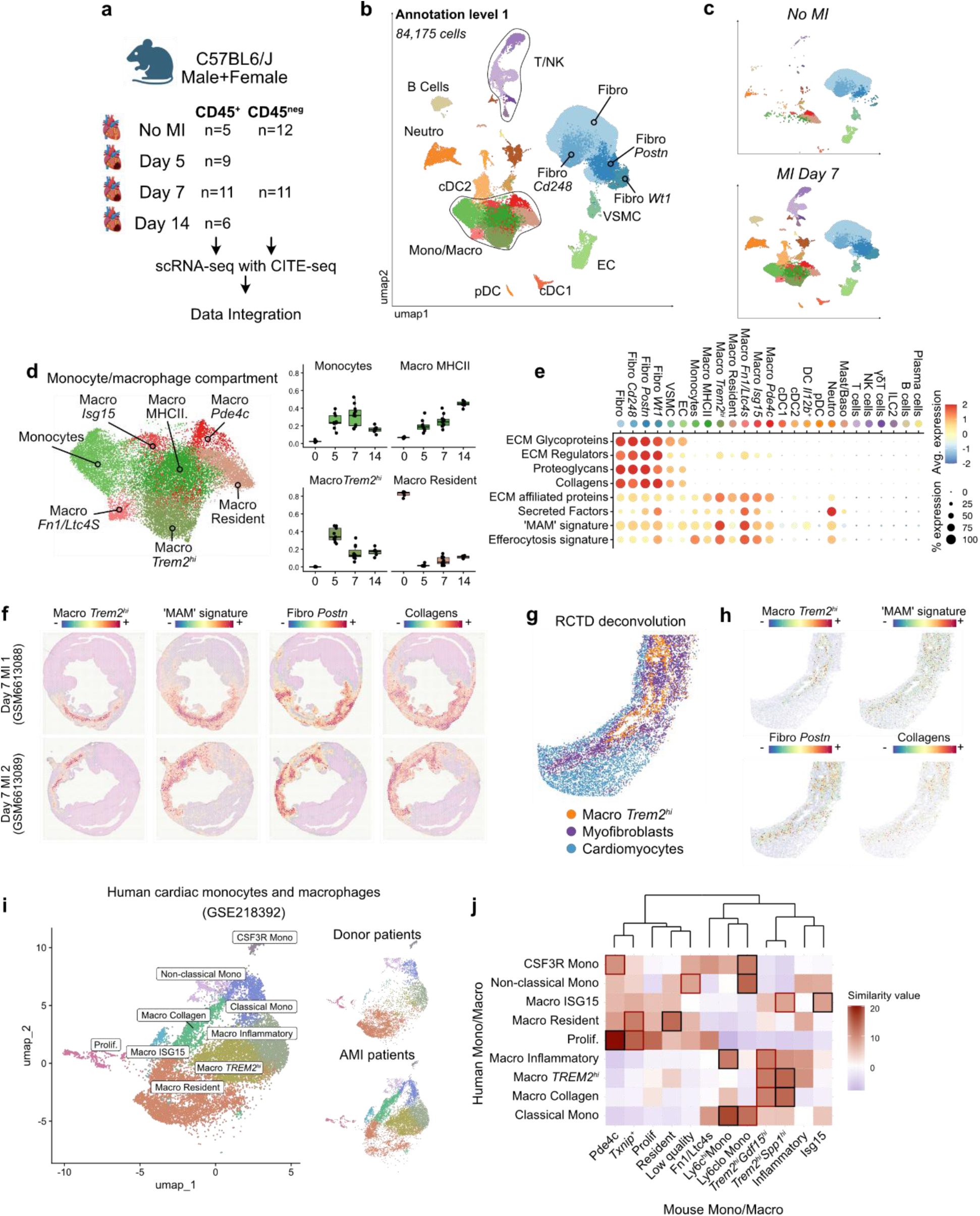
*Trem2^hi^* macrophages co-localize with myofibroblasts in the infarcted heart. **a)** Outline of the scRNA-seq experimental design. **b)** Uniform manifold approximation and projection (UMAP) plot of cardiac cells isolated from the infarcted heart at baseline (no MI), and at days 5, 7 and 14 post-MI. No MI n= 5 (CD45+) or n=12 (CD45-), MI Day 5 n= 9 (only CD45+), MI Day 7 n= 11 (CD45+ and CD45-), MI Day 14 n= 6 (only CD45+). **c)** UMAP of cardiac cells populating the heart at baseline (top) and at day 7 post-MI (bottom). **d)** UMAP plot of cardiac macrophages identified in b and re-clustered (left) and proportion of the main macrophage population at Baseline (0) and at days 5, 7 and 14 post-MI (right). **e)** DotPlot showing gene signatures score in total cardiac cells. Gene lists to calculate these scores were retrieved from the Matrisome Project (31) or from Gene Ontology term GO:0043277 ‘Apoptotic Cell Clearance’ for the efferocytosis score. **f)** Projection of the gene signatures onto Visium spatial transcriptomics data from day 7 post-MI hearts identifying the localization of the clusters ‘Macro Trem2’ and ‘Fibro Postn’, and ‘MAM’ and ‘Collagens’ signatures expression. **g)** Visualization of the indicated cell populations in the Curio Seeker spatial transcriptomics data, obtained from a day 7 post-MI heart. The predominant cell type in each spot was identified by performing deconvolution using the Robust Cell Type Decomposition (RCTD) tool (27). **h)** Spatial localization of the indicated cell populations (‘Macro Trem2’ and ‘Fibro Postn’) and gene signatures (‘MAM signature’ and ‘Collagens’) in the Curio Seeker spatial transcriptomics data. **i)** UMAP plot of cardiac monocytes and macrophages from donor patients and after acute myocardial infarction (AMI). Donor n=6, MI n= 4, data from GSE218392. **j)** Human-mouse cross species cluster similarity analysis of cardiac monocytes and macrophages represented as a Heatmap. Black highlight indicates top first similarity, while the red highlight indicates the top second similarity.

The ‘Macro *Trem2*’ cluster peaked at day 5 post-MI, coinciding with the onset of myofibroblast activation and scar formation (**Figure 1d**). Consistent with previous findings describing the role of *Trem2^hi^* macrophages in fibrotic processes, our data prompted us to hypothesize that cardiac *Trem2^hi^* macrophages support fibrosis in post-MI remodeling. To corroborate this hypothesis, we analyzed the expression of gene signatures extracted from the Matrisome Project (31) in our scRNA-seq dataset. Fibroblasts exhibited high expression of gene signatures corresponding to ‘extracellular matrix (ECM) glycoproteins’, ‘ECM regulators, ‘proteoglycans’, ‘collagens’, ‘ECM affiliated proteins’ and ‘secreted factors’ (**Figure 1e**). The ‘Macro *Trem2’* and ‘Macro *Fn1*/*Ltc4s*’ clusters also displayed elevated scores for ‘ECM affiliated proteins’ and ‘Secreted factors’ (**Figure 1e**). The ‘Macro *Trem2*’ cluster was also highly enriched in a matrisome-associated macrophage (MAM) signature, previously reported to be conserved among macrophages with pro-fibrotic function in different organ contexts (16). TREM2 is involved in phagocytosis processes, including efferocytosis (19), and genes associated with efferocytosis (Gene Ontology GO:0043277 ‘Apoptotic Cell Clearance’) were strongly expressed in the ‘Macro *Trem2*’ and ‘Macro *Fn1*/*Ltc4s*’ clusters (**Figure 1e**), suggesting a high efferocytic gene program in *Trem2^hi^* macrophages. Analysis of the monocyte/macrophage compartment with higher granularity revealed two distinct *Trem2^hi^* macrophage clusters with discrete signatures named *Trem2^hi^Spp1^hi^*and *Trem2^hi^Gdf15^hi^* (**Supplemental Figure 1B-D**), consistent with previous findings from our group, in which we identified *Trem2^hi^Spp1^hi^* as monocyte-to-macrophage intermediate state and the *Trem2^hi^Gdf15^hi^* as a fully differentiated population (3). Although expression of the MAM signature gene was elevated in both *Trem2^hi^* macrophage populations, the *Trem2^hi^Spp1^hi^* cluster showed an enhanced MAM signature score compared to the *Trem2^hi^Gdf15^hi^* cluster (**Supplemental Figure 1E**), suggesting a more specialized pro-fibrotic program within the *Trem2^hi^Spp1^hi^*cluster.

By reanalyzing publicly available spatial transcriptomics datasets of infarcted murine hearts (32), we confirmed that the “Macro *Trem2*” cluster and its associated MAM signature explicitly co-localize within regions enriched for *Postn^+^* myofibroblasts and collagen transcripts (**Figure 1f**), suggesting that *Trem2^hi^* macrophages may interact with fibroblasts. To further interrogate *Trem2^hi^* macrophage-fibroblasts interaction, we performed spatial transcriptomics of the scar and border zone at day 7 post-MI using the Curio Seeker Spatial Mapping Kit, allowing for whole-transcriptome spatial analysis with near single-cell resolution (**Figure 1g, Supplemental Figure 1F-H**). For cell type identification we applied Robust Cell Type Decomposition (RCTD) (27) onto the spatial transcriptomics data (**Supplementary Figure 1H**). This analysis revealed close interfacing between *Trem2^hi^*macrophages and myofibroblasts (**Figure 1g-h**).

To establish the translational relevance of our murine findings to human pathology, we re-analyzed publicly available scRNA-seq datasets from MI patients to comprehensively characterize their cardiac monocyte and macrophage landscapes (33) (**Figure 1i**). We identified a resident macrophage (‘Macro Resident’), two pro-inflammatory macrophages (‘Macro Inflammatory’ and ‘Macro ISG15’), two pro-fibrotic macrophage clusters (‘Macro TREM2^hi^’ and ‘Macro Collagen’), three monocyte clusters (‘Classical Mono’, ‘Non-classical Mono’ and ‘CSF3R Mono’) and a cluster of proliferating cells (‘Prolif.’) (**Figure 1i** and **Supplemental Figure 1I**). Similar to the mouse model, TREM2^hi^ macrophages were enriched in patients with MI compared to donor controls (**Supplemental Figure 1J**). Notably, an elevated MAM signature score was observed in the “Macro TREM2^hi^” cluster (**Supplemental Figure 1K**). To evaluate conserved macrophage responses across species, we conducted a transcriptional similarity analysis between our murine data (**Supplemental Figure 1B**) and the human datasets (**Figure 1I**) using the ClusterFoldSimilarity R package (28). This comparison revealed a high degree of similarity between human monocytes and macrophages and their murine counterparts. Notably, the human “Macro TREM2^hi^” cluster exhibited strong similarity to the mouse “Trem2_*Gdf15*” and “Trem2_*Spp1*” clusters (**Figure 1j and Table 1**), demonstrating a conserved pro-fibrotic response in *TREM2^hi^* macrophages in humans. In summary, we identified a conserved *Trem2^hi^* macrophage population that is highly enriched in pro-fibrotic and efferocytic genes, and spatially co-localizes with myofibroblasts within the infarcted myocardium.

### *Trem2* drives accumulation of monocyte-derived macrophages with a pro-scarring matrisome-associated gene expression profile in the infarcted heart

TREM2 is essential for the accumulation of lipid-associated macrophages in the obese adipose tissue (6). To assess the role of TREM2 in macrophage accumulation in the infarcted heart, we obtained tissue sections of hearts from *Trem2^+/+^* and *Trem2^-/-^* mice at day 5 post-MI, and quantified macrophage area by staining for CD68 (**Figure 2a**). Macrophage coverage was reduced in the infarcted area in *Trem2^-/-^* compared to *Trem2^+/+^*mice (**Figure 2a**). We then performed scRNA-seq/CITE-seq analysis of total CD45^+^ cells in *Trem2^+/+^* and *Trem2^-/-^* mice before and at 5 days after MI, to evaluate how TREM2 affects specific MI-associated macrophage subsets (3) (**Supplementary Figure 2A-B**). Monocytes/macrophages were identified and re-clustered to precisely characterize macrophage subsets (**Figure 2b**). We recovered macrophage clusters similar to those we previously identified (3) (**Figure 2b and Supplemental Figure 2C**), including *Trem2^hi^Spp1^hi^*and *Trem2^hi^Gdf15^hi^* macrophages. Although macrophages from *Trem2^-/-^* mice do not express *Trem2* (**Supplementary Figure 2D**), we still employ the ‘*Trem2^hi^*’ denomination for clarity. We observed preferential reduction of the *Trem2^hi^Spp1^hi^* cluster and Ly6C^hi^ monocytes in *Trem2^-/-^* mice compared to control (**Figure 2c**), indicating decreased accumulation of a major matrisome-associated monocyte-derived macrophage population associated with *Trem2* deficiency. The MAM and ‘ECM regulators’ gene expression scores were furthermore reduced in the *Trem2^hi^Spp1^hi^* and *Trem2^hi^Gdf15^hi^* clusters in *Trem2^-/-^* mice compared to control (**Figure 2d**). Gene expression scores for ‘Secreted factors’, ‘Proteoglycans’, and ‘ECM Glycoproteins’ were preferentially downregulated in the *Trem2^hi^Gdf15^hi^* subset in *Trem2^-/-^*mice, while expression of genes linked to apoptotic cell clearance and “ECM regulators” were decreased in the *Trem2^hi^Spp1^hi^* subset in *Trem2^-/-^* mice (**Figure 2d**). Altogether, our data indicate that TREM2 is involved in the accumulation of *Trem2*^hi^ monocyte-derived macrophages in the infarcted heart and modulates their matrisome-associated gene expression signature. Furthermore, TREM2 appears to differentially regulate the *Trem2^hi^Spp1^hi^* and *Trem2^hi^Gdf15^hi^* macrophage subsets, suggesting specific regulatory mechanisms within these populations.

**Figure 2.**
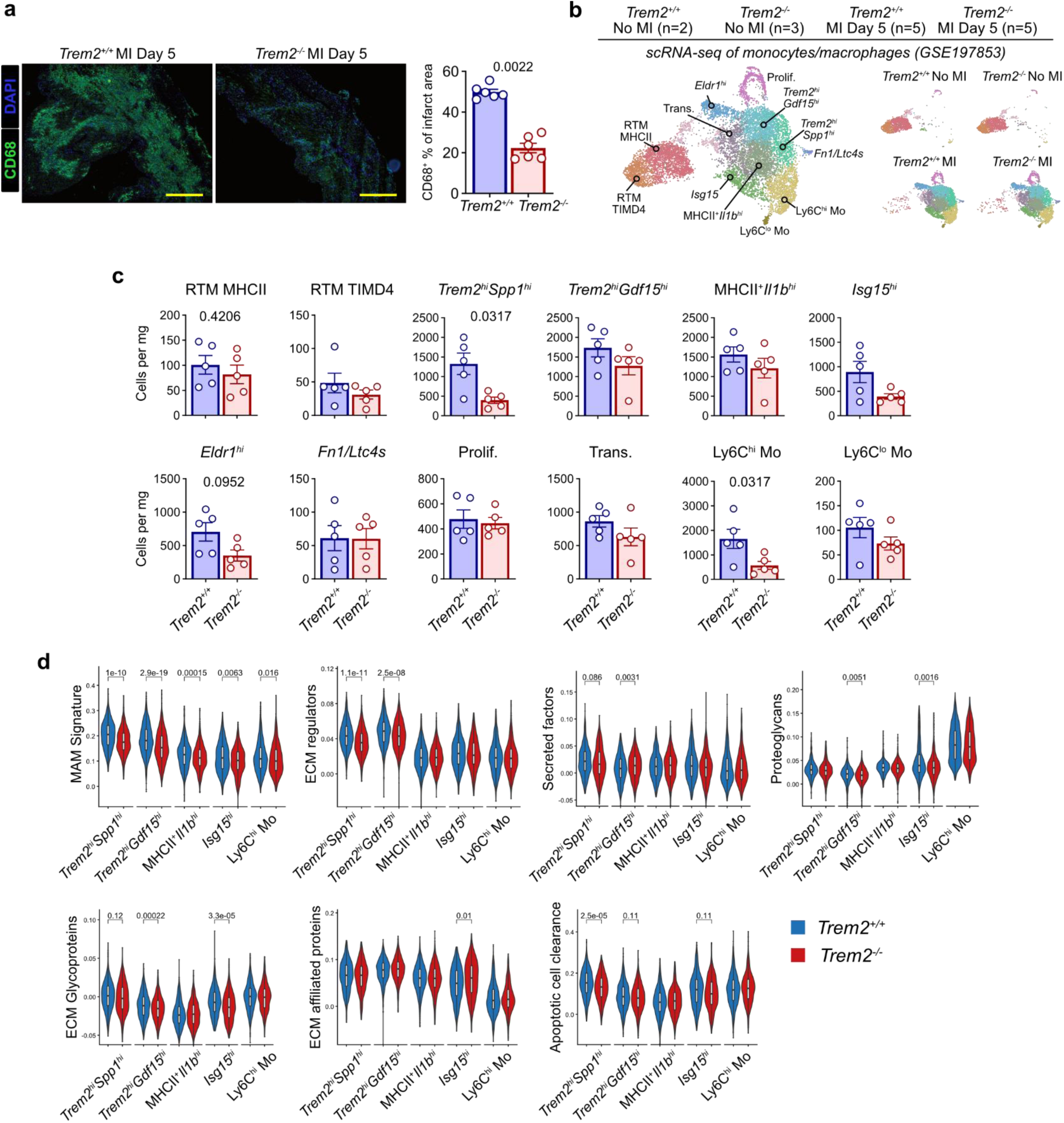
*Trem2* drives accumulation of monocyte-derived macrophages with a pro-scarring matrisome-associated gene expression profile in the infarcted heart. **a)** Representative pictures of CD68 immunofluorescence of cardiac tissue at days 5 post-MI (left) from *Trem2^+/+^* and *Trem2^-/-^* and quantification of CD68+ area (right). CD68 = green, DAPI = blue. *Trem2^+/+^* n= 6 and *Trem2^-/-^* n = 6. **b)** UMAP plot of annotated cardiac monocyte and macrophage clusters (left) and sample of origin (right). *Trem2^+/+^* No MI n= 2, *Trem2^-/-^* No MI n = 3, *Trem2^+/+^* MI Day 5 n= 6, *Trem2^-/-^* MI Day 5 n = 6. **c)** Cell counts per mg of cardiac tissue on the indicated cell populations in *Trem2^+/+^* and *Trem2^-/-^* mice. *Trem2^+/+^* MI Day 5 n= 6, *Trem2^-/-^* MI Day 5 n = 6. **d)** Expression of the indicated fibrotic scores in monocytes and macrophages clusters in *Trem2^+/+^* and *Trem2^-/-^*mice. *Trem2^+/+^* MI Day 5 n= 6, *Trem2^-/-^* MI Day 5 n = 6.

### *Trem2* deficiency impairs myofibroblast accumulation, collagen deposition and scar formation in post-MI cardiac repair

We then hypothesized that TREM2 is involved in the pro-fibrotic macrophage response by modulating fibroblasts. Immunofluorescence analysis revealed reduced αSMA^+^ myofibroblast coverage in *Trem2^-/-^*mice compared to *Trem2^+/+^* at day 7 post-MI (**Figure 3a**). In line, flow cytometric analysis revealed a reduced number of Ki67^+^ proliferating fibroblasts in infarcted *Trem2^-/-^* hearts compared to control (**Figure 3b**). Collectively, these observations demonstrate that TREM2 deficiency limits fibroblast proliferation and accumulation in the infarcted myocardium *in vivo*. To evaluate the chronic pathophysiological consequences of this impaired cellular response, we next characterized long-term cardiac remodeling up to 28 days post-MI. Survival was not affected in *Trem2^-/-^* mice compared to control (**Figure 3c**). A trend towards decreased ejection fraction was observed in *Trem2^-/-^* mice compared to control at day 7 after MI, but no differences at baseline, day 14 and day 28 after MI were noted (**Figure 3d**). Further histological analysis revealed increased infarct size in *Trem2^-/-^* mice compared to control (**Figure 3e**). Interestingly, collagen deposition in the infarct border zone was reduced in *Trem2^-/-^* compared to *Trem2^+/+^* mice (**Figure 3f**), while it was unchanged in the non-ischemic remote zone (**Figure 3g**). Angiogenesis and cardiomyocyte cross-sectional area were unchanged in *Trem2*^-/-^ mice compared to control (**Figure 3h-i**). Altogether, these results show that TREM2 is an important regulator of post-MI scarring and fibrotic processes mediated by macrophages, potentially by modulating fibroblast function and collagen deposition in the infarcted heart.

**Figure 3.**
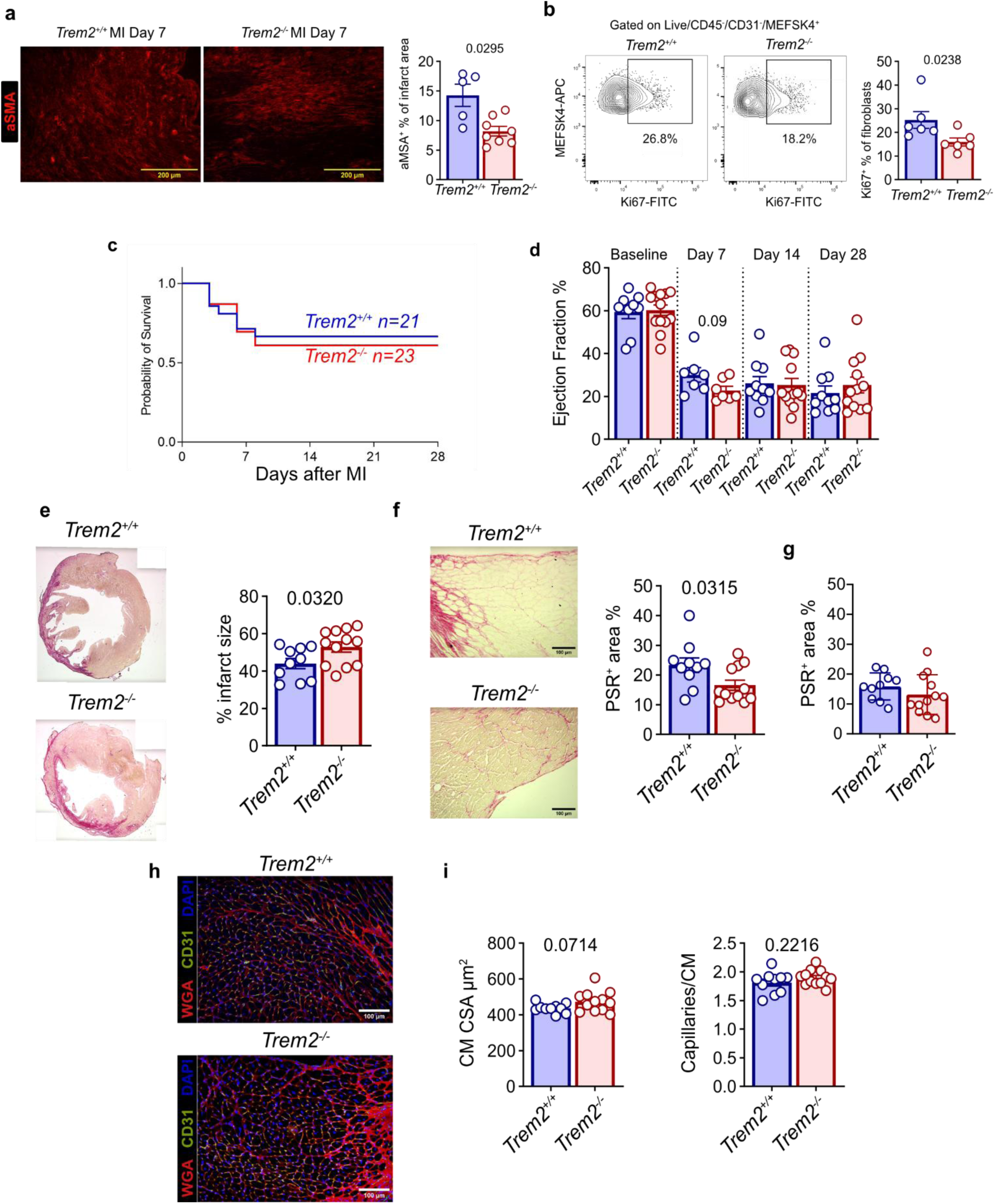
Trem2 deficiency impairs myofibroblast accumulation, collagen deposition and scar formation in post-MI cardiac repair. **a)** Representative heart pictures of αSMA (red) staining (left) and its quantification (right) at days 7 post-MI in *Trem2^+/+^*and *Trem2^-/-^* mice. *Trem2^+/+^* n= 5, *Trem2^-/-^* n = 8. **b)** Flow cytometry plots (left) of Ki67 staining in cardiac fibroblasts the heart and its quantification (right) at days 7 post-MI in *Trem2^+/+^* and *Trem2^-/-^* mice. *Trem2^+/+^* n= 6, *Trem2^-/-^* n = 6. **c)** Survival of males *Trem2^+/+^*and *Trem2^-/-^* mice over 28 days post-MI. *Trem2^+/+^*n= 21, *Trem2^-/-^* n = 23. **d)** Ejection fraction of males *Trem2^+/+^* and *Trem2^-/-^* mice measured at baseline, and at days 7, 14 and 28 post-MI. Baseline: *Trem2^+/+^* n= 9 and *Trem2^-/-^* n= 13, MI Day 7: *Trem2^+/+^* n= 7 and *Trem2^-/-^* n= 7, MI Day 14: *Trem2^+/+^* n= 10 and *Trem2^-/-^* n = 12, MI Day 28: *Trem2^+/+^* n= 10 and *Trem2^-/-^* n= 12. **e)** Representative picrosirius stained heart cryosection pictures (left) and infarct size quantification (right) in *Trem2^+/+^* and *Trem2^-/-^* mice at day 28 after MI. *Trem2^+/+^* n= 10 and *Trem2^-/-^* n = 12. **f)** Representative Picrosirius red staining of the heart border zone (left) and quantification of interstitial fibrosis (right) in *Trem2^+/+^* and *Trem2^-/-^* mice at day 28 after MI. *Trem2^+/+^* n= 10 and *Trem2^-/-^* n = 12; ; scalebar 100µm, magnification 20X. **g)** Picrosirius red staining quantification of the heart remote zone in *Trem2^+/+^* and *Trem2^-/-^*mice at day 28 after MI. *Trem2^+/+^* n= 10 and *Trem2^-/-^*n = 12. **h)** Representative immunofluorescence pictures of the heart border zone in *Trem2^+/+^* and *Trem2^-/-^* mice at day 28 after MI. Red = WGA, green = CD31, blue = Dapi. *Trem2^+/+^* n= 10 and *Trem2^-/-^* n = 12; scalebar 100µm, magnification 20X. **i)** Quantification of cardiomyocytes cross-sectional area (right) and capillaries per cardiomyocytes (left) in *Trem2^+/+^* and *Trem2^-/-^*mice at day 28 after MI. *Trem2^+/+^* n= 10 and *Trem2^-/-^*n = 12.

### Efferocytosis drives acquisition of the pro-scarring macrophage signature

In our previous studies, we pinpointed efferocytosis of apoptotic cells as a driver of *Trem2* and *Trem2*-related gene programs in bone marrow-derived macrophages (BMDM) (3), (19). In the infarcted heart, macrophages phagocyte necrotic cardiomyocytes (34) and apoptotic neutrophils (35), and we observed increased expression of genes associated with efferocytosis in *Trem2^hi^* macrophages (**Figure 1e**). In particular, engulfment of apoptotic neutrophils induces the shift from pro-inflammatory to anti-inflammatory response in macrophages (35). In addition, platelets have been shown to drive pro-fibrotic macrophage differentiation following MI (36). To determine which stimuli drives the pro-fibrotic program of macrophages recruited to the ischemic heart, we challenged BMDM with apoptotic neutrophils, necrotic cardiomyocytes or activated platelets to mimic the cardiac environment after MI (**Figure 4a**). Gene expression analysis revealed increased expression of fibrotic genes, such as *Timp2*, *Fn1*, *Mmp14*, *Tgfb1*, *Gpnmb* and *Spp1*, in BMDM exposed to apoptotic neutrophils, while no upregulation of these genes was observed in response to necrotic cardiomyocytes or activated platelets (**Figure 4a**). This is in line with our previous results showing acquisition of markers of the *Trem2* signature in response to apoptotic cell efferocytosis (3), (19).

**Figure 4.**
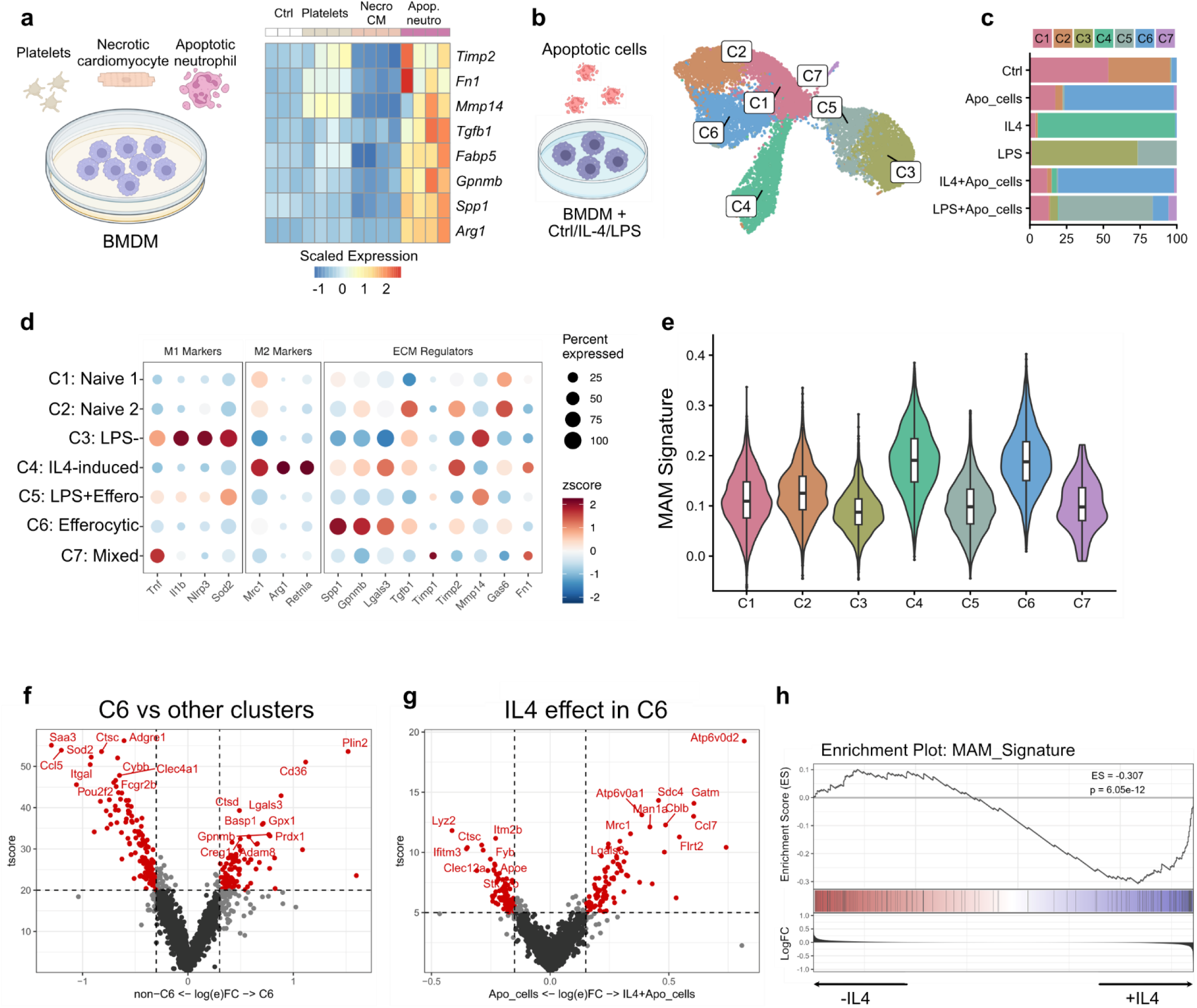
Efferocytosis drives acquisition of the pro-scarring macrophage signature. **a)** left: schematic representation of the experimental setup; right: Quantitative Real-Time Polymerase Chain Reaction (RT-qPCR) of the indicated genes in bone marrow-derived macrophages (BMDM) control (Ctrl), and treated with platelets, necrotic cardiomyocytes (Necro CM) and apoptotic neutrophils (Apop. neutro). Ctrl n= 4, Platelets n= 4, Necro CM n= 4 and Apop. neutro n= 4. **b)** Schematic representation of the experimental setup (left) and UMAP plot (right) showing the macrophage clusters identified from the scRNA-seq data. Ctrl n= 3, Apo_cells n= 3, IL4 n= 3, LPS n=2, IL4+Apo_cells n=3, LPS+Apo cells n= 2. **c)** Bar plot showing cluster distribution across experimental condition from the scRNA-seq data of **b**. Ctrl n= 3, Apo_cells n= 3, IL4 n= 3, LPS n=2, IL4+Apo_cells n=3, LPS+Apo cells n= 2. **d)** Dotplot showing the expression of M1, M2 and ECM regulators genes in the clusters identified in the scRNA-seq data of **b**. Ctrl n= 3, Apo_cells n= 3, IL4 n= 3, LPS n=2, IL4+Apo_cells n=3, LPS+Apo cells n= 2. **e)** Expression of the MAM signature genes in BMDM cluster across experimental conditions. Ctrl n= 3, Apo_cells n= 3, IL4 n= 3, LPS n=2, IL4+Apo_cells n=3, LPS+Apo cells n= 2. **f)** Differentially expressed genes identified between C6 against the other clusters (C1-C5). **g)** Differential gene expression analysis of cluster C6 between IL4+Apo_cells and Apo_cells conditions. **h)** GSEA enrichment score showing MAM score in C6 comparing Apo_cells with IL4+Apo_cells.

While IL-4 exposure is known to induce *Trem2^hi^* macrophage markers in BMDMs (37), recent work suggests that the synergy between type 2 cytokines and active efferocytosis is required to fully reprogram macrophages toward a pro-healing phenotype (38). To investigate whether efferocytosis-induced pro-fibrotic macrophage responses require IL-4 priming, we performed scRNA-seq on BMDMs stimulated with apoptotic Jurkat cells (as a standardized efferocytic bait), IL-4, or LPS, either alone or in combination with apoptotic cells (**Figure 4b**). Unbiased clustering revealed substantial heterogeneity among BMDMs across conditions, identifying 7 distinct clusters of BMDMs (C1 to C7). C1 and C2 were predominantly detected in untreated, baseline BMDMs. C3 was enriched in LPS-treated BMDMs, while C4 was mainly represented in IL-4-treated BMDMs. C5 was present in both LPS and LPS plus apoptotic cell-treated BMDMs. C6 represented the major population in macrophages stimulated with apoptotic cells alone or in combination with IL-4. Finally, C7 represented a minor cluster present in all the conditions (**Figure 4c**). ‘M1-like’ marker genes were predominantly expressed in C3, whereas ‘M2-like’ genes were highly enriched in C4, consistent with the classical M1/M2 macrophage polarization observed *in vitro* upon LPS and IL-4 stimulation (39) (**Figure 4d**). C6 showed strong enrichment in pro-fibrotic genes, including *Spp1*, *Gpnmb*, *Lgals3*, *Tgfb1*, *Timp2* and *Gas6*, as well as a high MAM signature score (**Figure 4d-e**). Differential gene expression analysis comparing cluster C6 against all the other clusters further confirmed the upregulation of pro-fibrotic genes, such as *Gpnmb*, *Lgals3* and *Prdx1* (**Figure 4f**). The MAM signature score was elevated in clusters C4 and C6, suggesting that IL-4 and efferocytosis are the main drivers of the pro-fibrotic response. To determine whether efferocytosis and IL-4 stimulate overlapping pro-fibrotic pathways, we evaluated the ‘extracellular matrix organization’ gene sets from the Reactome database across clusters C4 and C6. This analysis revealed a distinct, differential enrichment of matrix-remodeling pathways between these two cellular subsets (**Supplemental Figure 3C**). Moreover, pro-fibrotic genes such as *Gpnmb*, *Spp1*, and *Lgals3* were increased in C6 compared to C4. In contrast, *Timp2* expression was elevated in C4, while *Tgfb1* and *Timp1* remained unaffected (**Figure 4d and Supplemental Figure 3D**), indicating that efferocytosis and IL-4 induce distinct fibrotic transcriptional programs. Given the distinct roles of both efferocytosis and IL-4 in orchestrating macrophage fibrotic responses, we next investigated whether IL-4 priming could synergistically boost the pro-fibrotic program of efferocytic macrophages. Comparison of cells within the C6 in efferocytic conditions in the presence or absence of IL-4 further revealed upregulation of lysosomal genes (e.g. *Atp6v0a1* and *Atp6v0d2*), known to control extracellular matrix degradation (40), in the presence of IL-4 (**Figure 4g**). Additionally, GSEA analysis revealed increased expression of MAM signature genes in IL-4 primed efferocytic BMDM (**Figure 4j**). Collectively, these findings indicate that efferocytosis actively reprograms macrophages toward a pro-fibrotic transcriptomic profile, a response that is further amplified by IL-4 priming.

### TREM2-dependent efferocytosis drives macrophage mediated fibroblast activation

We show that efferocytosis is a major trigger of the macrophage pro-fibrotic response. However, it remains unclear how these macrophages exert their pro-fibrotic function and how it is regulated by TREM2. *In vivo*, we observed reduced gene expression scores of efferocytosis, ECM regulators and secreted factors in *Trem2^-/-^* macrophages in the infarcted heart compared to control (**Figure 2d**), while, *in vitro*, efferocytosis upregulated the expression of pro-fibrotic mediators, such as *Gpnmb*, *Tgfb1* and *Spp1* (**Figure 4a**). We therefore hypothesized that efferocytosis triggers the expression and release of pro-fibrotic factors in macrophages in a TREM2-dependent manner. To evaluate the role of TREM2 in the macrophage pro-fibrotic response, we analyzed the expression of pro-fibrotic markers in efferocytic *Trem2^-/-^* BMDMs. The expression of *Gpnmb* and *Tgfb1* was markedly reduced, while *Spp1* expression was elevated, and *Mmp14*, *Timp2*, and *Fn1* levels remained unchanged compared to *Trem2^+/+^* cells (**Figure 5a**). To determine if this TREM2-dependent response modulates fibroblast activation, we stimulated fibroblasts with conditioned medium from efferocytic *Trem2^-/-^*and *Trem2^+/+^* BMDM. Conditioned medium from efferocytic *Trem2^+/+^*BMDMs significantly enhanced fibroblast migration compared to medium from non-efferocytic BMDMs. This effect was completely abolished when fibroblasts were treated with conditioned medium from efferocytic *Trem2^-/-^* BMDMs (**Figure 5b-c**). These results indicate that TREM2 is required for the production of pro-fibrotic mediators following efferocytosis. Among these pro-fibrotic factors, *Gpnmb* expression was the most affected in *Trem2^-/-^*BMDMs and its expression was also reduced in *Trem2^hi^* macrophages *in vivo* in *Trem2^-/-^* mice compared to control (**Figure 5a and Figure 5d**). Furthermore, the GPNMB receptors Ryk and Gpr39 (41) (42) were specifically expressed in cardiac fibroblasts (**Figure 5e**), suggesting that the GPNMB-RYK-GPR39 axis could be responsible for the pro-fibrotic function mediated by *Trem2^hi^* macrophages. Altogether, our results place efferocytosis as the main driver of the TREM2-mediated pro-fibrotic macrophage response, supporting the upregulation of pro-fibrotic mediators, such as GPNMB.

**Figure 5.**
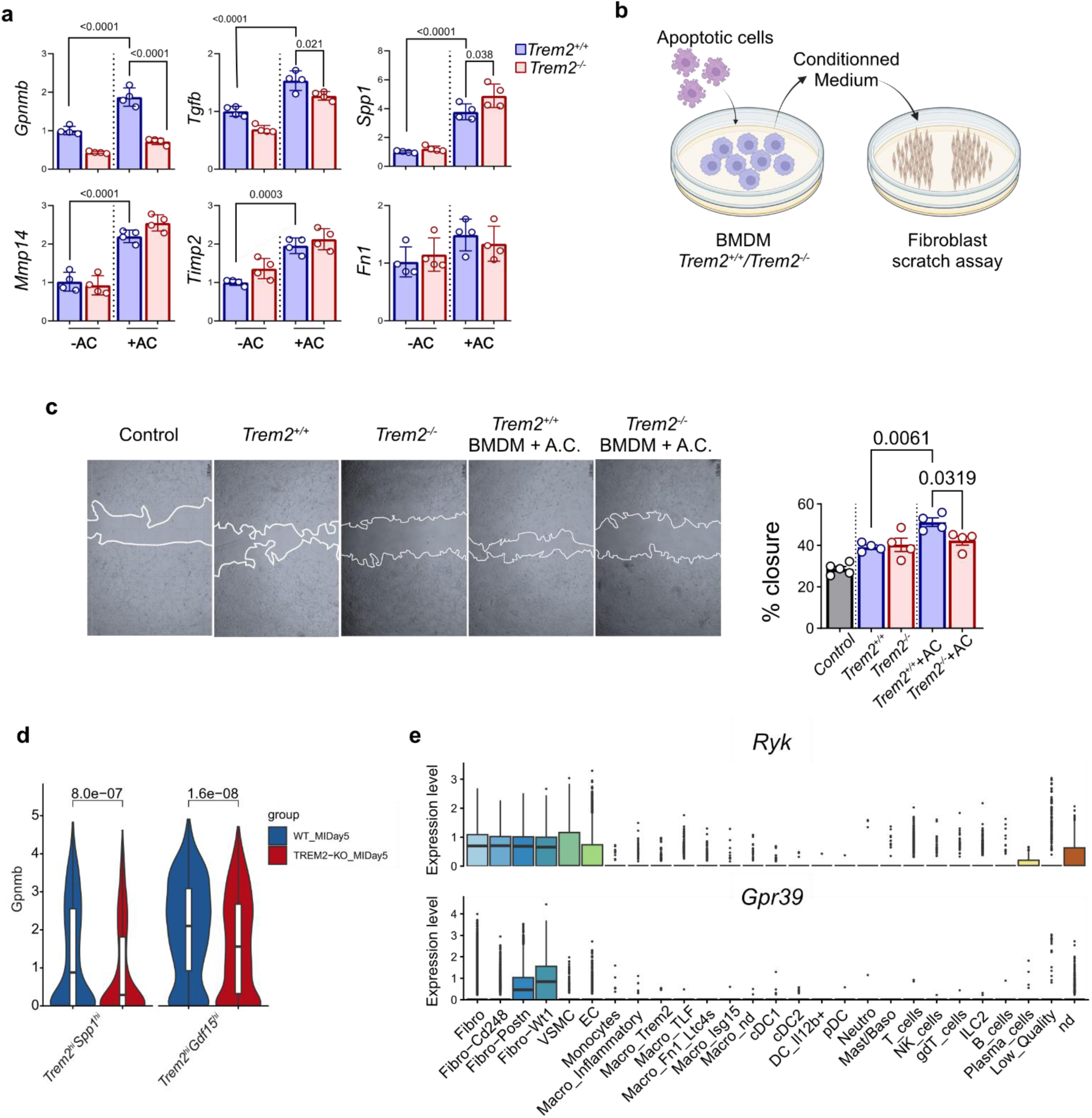
TREM2-dependent efferocytosis drives macrophage mediated fibroblast activation. **a)** qPCR analysis of the indicated genes in *Trem2^+/+^* and *Trem2^-/-^*BMDM under efferocytic challenge. *Trem2^+/+^* -AC n= 4, *Trem2^-/-^*-AC n= 4, *Trem2^+/+^* +AC n= 4, *Trem2^-/-^* +AC n= 4. **b)** Schematic representation of the scratch assay’s experimental setup. **c)** representative microscopic pictures of the scratch assay (left) and quantification of wound closure (right) 12 hours after stimulation. Control n= 4, *Trem2^+/+^*n= 4, *Trem2^-/-^* n= 4, *Trem2^+/+^*+AC n= 4, *Trem2^-/-^*+AC n= 4. **d)** *Gpnmb* expression in *Trem2^hi^* macrophages from the data in Figure 2b in *Trem2^+/+^*and *Trem2^-/-^* cells. **e)** Violin plot showing the expression of the indicated transcript in scRNA-seq data from Figure 1b.

## Discussion

In this study, we sought to elucidate the precise role of TREM2 in modulating macrophage function during post-infarction cardiac remodeling. Consistent with prior reports from our group and others, we observed a robust accumulation of *Trem2^hi^* macrophages within the infarcted myocardium during the acute inflammatory phase. Such *Trem2^hi^*macrophage subsets have been widely documented across diverse pathological settings, including neurodegenerative, metabolic, and fibrotic diseases (6), (9), (3), (13). Importantly, this *Trem2^hi^* macrophage population is highly conserved across species, expanding in both murine and human infarcted hearts, where we previously established their origin from recruited blood monocytes (3). While experimental evidence positions TREM2 as a highly promising therapeutic target with a proven beneficial role in cardiac remodeling, its underlying molecular and cellular mechanisms have remained elusive. Our study directly addresses this gap by deciphering how TREM2 mechanistically orchestrates the post-MI microenvironment.

By integrating single-cell and spatial transcriptomics, we demonstrated that cardiac *Trem2^hi^* macrophages peak at day 5 post-MI, harbor a distinct matrisome-associated macrophage signature, and preferentially localize within the myofibroblast niche. This tight spatial proximity strongly implies a functional cellular crosstalk that drives scar formation and fibrotic remodeling. Myofibroblasts coverage and proliferation were reduced in *Trem2^-/-^*mice compared to control, suggesting that TREM2 plays a critical role in fibrosis.

Although fibrosis is considered detrimental in most contexts, after MI the formation of a fibrotic scar maintains organ structure and protects the heart from rupture. However, interstitial fibrosis in the non-injured area and in the infarct border zone is detrimental, as it favors the transition to heart failure or the occurrence of arrhythmias (43).

Here, *Trem2^-/-^* mice exhibited exacerbated infarct sizes alongside a concomitant reduction in border-zone interstitial fibrosis. These divergent findings unveil a dual, compartmentalized role for TREM2: while it critically supports reparative scar formation within the core, it simultaneously drives deleterious interstitial fibrosis in the surrounding tissue. This internal biological trade-off likely explains why the net impact on global ventricular function remained neutral in our model. Previous studies have identified TREM2 as a critical regulator of pro-fibrotic macrophage function across various pathologies. In experimental models of NASH, lung and renal fibrosis, TREM2 has been described to promote fibrosis by activating fibroblasts (9), (44), consistent with our findings. Conversely, Ganguly et al. showed that *Trem2^hi^* macrophages play an anti-fibrotic role in MASH regression by facilitating collagen degradation (15). These divergent outcomes underscore that the pro-versus anti-fibrotic drive of *Trem2^hi^* macrophages is highly context- and organ-dependent. Within the post-MI microenvironment, this functional dichotomy may be further dictated by spatial segragation. While the infarct core is predominantly dominated by monocyte-derived macrophages, the border zone retains a substantial population of embryonically derived tissue-resident macrophages (45). Consequently, TREM2 signaling may exert distinct, cell-type-specific effects depending on the ontogeny of the underlying macrophage subset.

In the infarcted heart, the absence of TREM2 significantly affected the pro-fibrotic macrophage response, with reduced accumulation of *Trem2^hi^*macrophages and expression of the pro-fibrotic matrisome-associated signature. These results are consistent with reduced interstitial fibrosis in *Trem2*^-/-^ mice, suggesting that TREM2 has a critical role in regulating the pro-fibrotic function of macrophages. This *in vivo* observation was further supported by our *in vitro* data, which identified efferocytosis as the primary trigger for the pro-fibrotic signature in BMDMs.

Among the targets regulated by efferocytosis, expression of the pro-fibrotic factors *Gpnmb* and *Tgfb1* was significantly blunted in *Trem2^-/-^* BMDM, suggesting that TREM2 critically modulates the macrophage secretome. Consistently, *Trem2^hi^* macrophages from *Trem2^-/-^*mice exhibited a significantly reduced ‘secreted factors’ module score, establishing the translational relevance of our *in vitro* findings to the *in vivo* setting.

Among these soluble mediators, *Gpnmb* was drastically affected in the absence of *Trem2* both *in vivo* and *in vitro*. Macrophage-derived GPNMB has been previously documented to orchestrate tissue repair and fibrogenesis in both the heart and liver. Mechanistically, it binds to its cognate receptors, RYK and GPR39, on fibroblasts, thereby triggering their activation and driving downstream proliferative responses (41), (42). Supporting this paracrine axis, conditioned medium from *Trem2^-/-^* efferocytic macrophages failed to stimulate fibroblast migration compared to control counterparts, providing functional evidence that TREM2 dictates the macrophage pro-fibrotic secretome. While Chan et al. established that IL-4 priming skews BMDMs toward a pro-repair phenotype (46), we expand this paradigm by demonstrating that IL-4 and efferocytosis act synergistically to drive a robust matrisome-associated signature. Consequently, we propose that IL-4 availability within the post-MI microenvironment represents a critical checkpoint that licenses and amplifies the pro-scarring potential of efferocytic macrophages.

To summarize, we propose *Trem2^hi^* monocyte-derived macrophages as critical mediators of cardiac repair post-MI, by promoting scarring and fibrosis through interaction with myofibroblasts. Mechanistically, efferocytosis of apoptotic cells, in combination with IL-4 availability, triggers a TREM2-dependent transcriptional program and the release of pro-fibrotic factors which are instrumental for fibroblast activation and migration.

## Acknowledgements

We thank the Single-Cell Center Würzburg for assistance with scRNA-seq experiments.

