## Supplemental material for "TREM2 drives accumulation of pro-scarring monocyte-derived macrophages in the infarcted myocardium"

**Supplemental Methods**

**Mice**

*Trem2*$^{-/-}$ mice were initially provided by Marco Colonna (Washington, USA) and bred in the ZEMM (Centre for Experimental and Molecular Medicine, Wuerzburg, Germany). Mice were exposed to 12 hours light cycle with lights turning on at 7:00 and switched off at 19:00. All animal studies conform to the Directive 2010/63/EU of the European Parliament and have been approved by the appropriate local authorities (Regierung von Unterfranken, Wuerzburg, Germany, Akt.-Z. 55.2-DMS-2532-2-743, Akt.-Z. 55.2.2-2532.2-865, Akt.-Z. 55.2.2-2532-2-1594-13).

**Myocardial infarction**

Aged matched 8- to 12-week-old *Trem2*$^{+/+}$ and *Trem2*$^{-/-}$ male mice on a C57BL6/J background underwent permanent ligation of the Left Anterior Descending (LAD) coronary artery to induce myocardial infarction. Briefly, 30 minutes prior surgery mice received an injection of analgesic (Buprenorphine, 0.1mg/kg, s.c.). Deep anesthesia was induced with 5% isoflurane and mice were intubated with endotracheal canula (B/Braun, 4269098S-01) and placed under mechanical ventilation (VentElite, Harvard Apparatus) on a heating pad (Physitemp Instruments inc., TCAT-2LV). After checking deep anesthesia by absence of paw withdrawal reflex, anesthesia was maintained by 2.0-2.5% during surgery. The heart was exposed by thoracotomy of the fourth intercostal space, the pericardium was removed, and LAD ligation was performed with a 7/0 non-resorbable nylon suture thread (Serapren, CP05341A). The rib cage was closed with two independent sutures and skin was closed with a continuous suture using a 6/0 non-resorbable nylon suture thread (Serapren, CP07281A). For pain alleviation, mice were injected twice daily with Buprenorphine (0.1mg/kg, s.c.) for two days after surgery.

**Echocardiography**

Echocardiography was performed using a Vevo 2100 System (FUJIFILM Visualsonics) and a MS400 transducer (FUJIFILM Visualsonics). C57BL6/J and *Trem2*$^{-/-}$ mice were kept in light anesthesia with 1% of Isoflurane, and echocardiography was recorded at day 0, 7, 14 and 28 after permanent coronary artery ligation. Long and short axis of the heart were acquired at apical, middle and base levels of the left ventricleM-mode of short-axis at the middle and apical levels were used to calculate ejection fraction. Only animals with a heart rate (HR) between 450 and 600 b/m were included in the analysis. Analysis was performed using Vevo LAB software version 3.2.6 (FUJIFILM Visualsonics).

**Histology**

*Trem2*$^{+/+}$ and *Trem2*$^{-/-}$ male mice were killed by cervical dislocation under isoflurane anesthesia and the hearts were injected with KCl before perfusing with ice-cold PBS. The heart was snap-frozen in liquid nitrogen-cold isopentane (Carl Roth, 3927.1). 7µm cryosections were made with a Cryostat (Leica, CM3050 S) onto SuperFrost slide (Langenbrick GmbH, 03-0060). For immunofluorescence stainings, heart sections were blocked with Normal Serum Block (Biolegend, 927503) for 30 minutes before primary antibody incubation with rat anti-mouse CD68 (Biolegend, clone FA-11, 137001, 2,5 µg/ml), rat anti-mouse CD31 (Invitrogen, clone 390, 14-0311-85, 1 µg/ml), Wheat Germ Agglutinin (Vector laboratories, RL-1022-5, 5 µg/ml), anti-αSmooth-muscle actin (Sigma-Aldrich, clone 1A4, C6198, 7 µg/ml) overnight. Then, heart sections were incubated with goat α-rat Alexa 488 (Invitrogen, A11006, 4 µg/ml). Finally, slides were mounted with Vectashield containing DAPI (Vectashield, H-1200). For picrosirius red staining, sections were passed in Xylene (VWR, 28975360) and decreasing concentrations of ethanol (CARL-ROTH, T913.3). After rinsing the sections twice with distilled water, the sections were incubated with Picrosirius red (Sigma, Direct Red 80, 365548-5G) for 1 hour. Following 2 passages in acidified water (0.5% glacial Acetic Acid (Sigma-Aldrich, 33209) in water), the sections were dehydrated in increasing ethanol concentrations and xylene. The slides were mounted with Vectashield (Vectashield, H-5000). Images were acquired with a Leica DM 4000 B LED microscope. Analysis was performed with ImageJ software (Fiji).

**Apoptotic cell generation**

Neutrophils were isolated from bone marrow cells of male C57BL6/J mice. Briefly, the tibia and femur were dissected to expose the bone marrow, which was then collected by placing them in a 0.5ml Eppendorf tube with a hole made at the bottom using a 12G needle. Subsequently, the 0.5ml Eppendorf tube containing the bones was inserted into a 1.5ml Eppendorf tube and centrifuged at 10000g for 15 seconds, allowing the bone marrow to be collected in the 1.5ml Eppendorf tube. The pellet was resuspended with MACS buffer (PBS) and neutrophils were isolated with anti-Ly6G Microbeads following manufacturer´s instruction (Miltenyi Biotech, 130-120-337). Apoptosis was induced by stimulating $1X10^6$ neutrophils/ml with 1µM of Staurosporine (Sigma Aldrich, S6942) in complete medium (RPMI (Gibco, 21875-034) supplemented with 10% FCS, 100 U/ml penicillin and streptomycin (Sigma-Aldrich, P4333), and 50µM of β-Mercaptoethanol (Gibco, 31350-010)) for 17 hours at 37°C. After washing twice, apoptosis was evaluated with Annexin V/7-AAD staining kit (ThermoFisher, 00-0055-56). Preparation with more more than 80% of Annexin V+7AAD- cell were used (**Supplemental Fig 3A**).

Apoptotic Jurkat T cells were generated by exposing Jurkat T cells to UV irradiation (312 nm) for 15 minutes in a 10-cm cell culture dish, followed by 12 hours rest in complete medium. Apoptosis was verified using the Annexin V/7-AAD staining kit (**Supplemental Fig 3B**).

**Necrotic cardiomyocytes generation**

Cardiomyocytes were isolated as described in (1). After isolation, cardiomyocytes were placed in starving medium (RPMI (Gibco, 21875-034) supplemented with 0,5% FCS, 100 U/ml penicillin and streptomycin (Sigma-Aldrich, P4333), and 50µM of β-Mercaptoethanol(Gibco, 31350-010)) and necrosis was induced by freeze/thawing at -20 °C.

**Platelet isolation**

Mouse platelets were isolated as previously described (2). Briefly, C56BL6/J mice were bled retro-orbitally under deep anesthesia in Heparin (Ratiopharm, Heparin- Natrium-5000, 5394.00.00). After two sequential centrifugations, plasma with no residual red blood cells was collected and centrifuged again to produce the platelet rich plasma (PRP). PRP was lastly washed again, and the resulting platelet pellet was resuspended in sterile PBS. Platelet count was measured with Sysmex (Automated Hematology Analyzer XP-300), and concentration was adjusted to $0.5X10^6$ platelets/µl. Platelet activation was induced with Thrombin (0,01U/ml, Merck/Roche 10602400001) for 5 minutes at 37°C.

**Bone marrow derived-macrophages (BMDMs) culture**

Bone marrow cells were extracted from male C57BL6/J and $TREM2^{-/-}$ mice. Tibia and femur were dissected to expose the bone marrow, which was then collected by placing them in a 0.5ml Eppendorf tube with a hole made at the bottom using a 12G needle. Subsequently, the 0.5ml Eppendorf tube containing the bones was inserted into a 1.5ml Eppendorf tube and centrifuged at 10000g for 15 seconds, allowing the bone marrow to be collected in the 1.5ml Eppendorf tube. After bone marrow isolation, the cell pellet was resuspended in complete medium (RPMI (Gibco, 21875-034) supplemented with 10% FCS, 100 U/ml penicillin and streptomycin (Sigma-Aldrich, P4333), and 50µM of β-Mercaptoethanol(Gibco, 31350-010)) supplemented with 15% L929 conditioned medium. Cells were counted and $2 \times 10^6$ cells per milliliter were plated in a 10-$cm^2$ cell culture dish. On day 7, adherent macrophages were harvested using Accutase (Sigma-Aldrich, A6964), washed, resuspended in complete medium, and $0.4X10^6$ macrophages were seeded into a 12-well plate (SARSTEDT, 83.3921). The following day, the macrophages were treated with starving medium. After a 4-hour resting period, the macrophages were stimulated with apoptotic neutrophils at a ratio of 1:5 (1 macrophage to 5 apoptotic neutrophils), necrotic cardiomyocytes at a ratio 5:1 (5 macrophages to 1 necrotic cardiomyocyte) and activated platelets at a 1:50 ratio (1 macrophage to 50 platelets) for 17 hours. BMDMs were then washed with PBS and resuspended in RA1 lysis buffer from the Nucleo Spin RNA extraction kit (Macherey-Nagel, 740955.50).

**Quantitative Real-Time Polymerase Chain Reaction**

RNA was isolated according to manufacturer´s instruction (Macherey-Nagel, 740955.50). Equal amount of RNA was converted in cDNA using random hexamer primer of the First Strand cDNA Synthesis kit (Thermofisher, K1612). Quantitative Real-Time Polymerase Chain Reaction (qPCR) was performed using SYBER-green (Applied Biosystems, A25742) and acquired on an Applied Biosystems QuantStudio 6 Fles Real-Time PCR System. The fold-change was calculated using the ΔCT method with Hprt as the housekeeping gene. Primer sequences are listed in **Table 2.**

**Scratch assay**

3T3 fibroblasts were plated in a 12 well plate and let them reach 80% confluence in fibroblast medium (DMEM (Sigma-Aldrich, D6429), 10% FCS and 100 U/ml penicillin and streptomycin (Sigma-Aldrich, P4333). Once confluent, they were placed in starving medium for 4 hours followed by the scratch with the help of a 1000 µl tip. The starving medium was then replaced by supernatant of macrophages treated with apoptotic Jurkat cells or control untreated. Images were acquired at 0 and 12 hours after stimulation using Nikon DS-L4 and analyzed with ImageJ (Fiji).

**Single-cell RNA-seq analysis**

<u>Generation of the non-myocyte cell atlas in mouse myocardial infarction (Figure 1)</u>

ScRNA-seq of viable immune cells ($CD45^{pos}$) and non-myocyte cells from non-infarcted and infarcted (day 5, 7 and 14 post-MI) male and female C57BL6/J mouse hearts were obtained from several independent experiments. In all experiments except the day 14 time point, mice were injected i.v. with 2.5µg anti-CD45.2 APC or PE (clone 104, Biolegend, cat. #109814) under isoflurane anesthesia 5 minutes before euthanasia to exclude intravascular cells during sorting as described in (3) and (4). The hearts were enzymatically digested at 37°C with agitation at 800 rpm/min in RPMI containing 450U/ml collagenase I (Sigma-Aldrich C0130), 125U/ml collagenase XI (Sigma-Aldrich C7657), 60U/ml Hyaluronidase (Sigma-Aldrich H3506), 60U/ml DNAse (Roche #11284932001) for 45 minutes, and target cells pre-enriched using either CD45+ magnetic sorting (Miltenyi Biotec #130-052-301) or magnetic Dead Cell Removal kit (Milteny Biotec #130-090-101) with LS-Columns (Milteny Biotec #130-042-901). In each experiment, biological replicates (i.e. cells from each individual mouse) were labeled with hashtag antibodies (BioLegend TotalSeq-A), pooled and processed together for cell sorting and scRNA-seq library generation. These experiments comprise: 2 experiments with sorting of immune cells (total viable $CD45^{pos}$ cells) from non-infarcted and day 5 post-MI hearts (GSE197441, GSE197853), 1 experiment with sorting of viable non-immune cells from non-infarcted hearts (viable $CD45^{neg}$, GSE328819; GSM9690204), 1 experiment with sorting of viable non-immune cells from day 7 infarcts (viable $CD45^{neg}$, GSE328819; GSM9690223), 1 experiment with sorting of viable immune cells from day 7 infarcts (viable $CD45^{pos}$, GSE328819; GSM9690219), and 1 experiment with sorting of viable immune cells from day 14 infarcts (viable $CD45^{pos}$). ScRNA-seq libraries were generated by using 10X Chromium Next GEM Single Cell 3′ Kit and 10X Chromium Next GEM Single Cell 3′ HT Kit version (10x Genomics) according to the manufacturer's instructions. Sequencing was performed using a NovaSeq6000. The 10x Genomics single-cell gene expression, HTO, ADT libraries were processed using Cell Ranger software (version 7.0.1) and the sequence alignment was performed with Mouse mm10 reference genome. scRNA-seq data analysis. Cell Ranger outputs were obtained in the form of filtered_feature_bc_matrix files and were loaded in R (version 4.4.2.) and further analyzed using Seurat v5 (5). For each experiment, data were pre-processed to assign each cell to its sample of origin based on the hashtag signal using the 'HTODemux' function in Seurat. Multiplets (i.e. cells with multiple hashtag signals) and cells without hashtag assignment were excluded from downstream analyses. Quality control thresholds were applied to remove cells with high mitochondrial RNA content (>5; >7.5 or >10% depending on experiments), or cells with outlier UMI counts likely corresponding to remaining multiplets. Cells from the relevant samples (i.e. infarcted or non-infarcted C57BL6/J mice without treatment or genetic modifications) were selected for downstream analyses. Pre-processed data from all experiments were pooled and batch corrected using the Harmony method within Seurat, using the 'IntegrateLayers' function with 'method = HarmonyIntegration'. Clustering and dimensional reduction were performed using 30 principal components and a resolution parameter of 1.0. Cell lineages and subpopulations were identified based on the expression of RNA and surface markers based on previous publications notably (4) for immune cells and (6) for non-immune cells and fibroblast subsets. To analyze macrophages with higher granularity, cells corresponding to monocytes and macrophages were extracted and re-clustered using 30 principal components and a 0.6 resolution, and annotated based on (4).

Analysis of WT and *Trem2*$^{-/-}$ cardiac immune cells at day 5 post-MI

We analyzed previously published data available in gene expression omnibus (GSE197853). Generation of the single-cell RNA-seq data from CD45$^{+}$ cells of infarcted (day 5) and non-infarcted Trem2$^{+/+}$ and Trem2$^{-/-}$ hearts was described in details in (4). Quality control of the two libraries was performed independently using the same pipeline. Count matrices were analyzed using Seurat v5 (5). Hashtag and CITE-seq antibody counts were separated from the main gene expression (RNA) matrix. RNA data was normalized using the NormalizeData function, the top 2,000 highly variable genes were identified and then scaled using the ScaleData function. Hashtag and CITE-seq data were added as independent assays and normalized using the NormalizeData with the centered log-ratio (CLR) transformation. Demultiplexing was performed to identify sample of origin, exclude multiplets, and cells with undetectable hashtag signal using the HTODemux function setting "positive.quantile = 0.9999" to keep only singlets. Dead cells and potential outliers were filtered out by excluding cells with more than 7.5% of mitochondrial genes and 40000 total RNA counts, respectively. After quality control, the two libraries were batch corrected with Harmony and sample identities were assigned based on the hashtag signal. Total $CD45^+$ cells clustering was performed with the FindCluster function in Seurat using 20 principal components and 0.6 resolution was used. The main immune cell population were identified using canonical markers as described previously (4). Monocytes and macrophages were identified in the total leukocyte pool and extracted for in depth analysis. Clustering analysis was performed using 20 principal components and a 0.8 resolution. Monocytes and macrophages were annotated based on the top30 marker genes identified with the FindAllMarkers function in Seurat. Monocyte and macrophage absolute counts were calculated by multiplying the cluster proportion with the absolute counts of $CD45^+$ cells per mg of tissue as determined using flow cytometry of the same samples with counting beads (see (4)). The Matrisome-associated macrophage (MAM) gene list signature was retrieved from published data (7) and applied using the AddModuleScore function to the monocyte and macrophage clusters. Signatures of ECM regulators, Secreted factors, Proteoglycans, ECM Glycoprotein, ECM affiliated proteins were retrieved from the Matrisome Project (https://sites.google.com/uic.edu/matrisome/home) and applied using the AddModuleScore function to the monocyte and macrophage clusters. The Apoptotic cell clearance signature was retrieved from Gene Ontology GO:0043277 'Apoptotic Cell Clearance'. To account for the lack of expression of *Trem2* in *Trem2*$^{-/-}$ mice, the *Trem2* gene was removed from gene lists used to apply signatures where appropriate. Statistical analysis was performed with a Wilcoxon rank-sum test using the VlnPlot2 function of the SeuratExtend package (8) specifying the argument "stat.method = "wilcox_test".

Analysis of BMDMs after efferocytosis and pre-stimulation with IL4 and LPS (Figure 4)

BMDMs were generated from male C57BL6/J mice as described above (see Apoptotic cell generation). After 7 days, BMDMs were detached with Accutase and $0.4X10^6$ macrophages were seeded in 12-well plate, and left to rest overnight. After 4 hours in starving medium (RPMI supplemented with 0.5% FCS), macrophages were pre-stimulated with 20 ng/ml of IL4 or 10 ng/ml of LPS for 4 hours. The stimulations were then removed and apoptotic Jurkat cells were added at a ratio of 1:5 (1 macrophage to 5 Jurkat cells) for 17 hours. In the only IL-4 and LPS conditions, IL-4 and LPS were additionally left for 17 hours. The cells were washed twice with ice-cold PBS and detached using Accutase. The following steps were performed at 4°C. Macrophages were incubated with TruStain FcX for 10 minutes to block unspecific antibody binding. After washing, each sample was incubated with a specific hashtag antibody (BioLegend TotalSeqA, 1:200) for 30 minutes. **Table 1** indicates the sample/hashtag correspondence. Cells were washed twice, pooled and washed once again. The pooled cell suspension was incubated with F4/80 PE-Cy7 (Biolegend 123114, 2 µg/ml), a mix of TotalSeq-A antibodies against surface markers (**Table 3**) and Fixable Viability Dye eFluor780 (1:1000; ThermoFisher, 65-0865-14) for 20 minutes. Macrophages were washed once, resuspended in PBS/1%FCS and viable $F4/80^+$ cells were sorted using a BD FACS Aria III (BD Biosciences) with a 100µm nozzle. Sorted cells were loaded in the 10X Genomics Chromium platform aiming for a capture target of 20,000 cells. scRNA-seq libraries were prepared using the 10x Genomics Chromium Next GEM Single Cell 3´kit v3 (10x Genomics) and processed following the manufacturer´s instructions. Sequencing was performed using the NovaSeq 6000 (Illumina) platform. 10x Genomics data, HTO and ADT libraries were demultiplexed using Cell Ranger software (version 7.0.1). Alignment and counting steps were performed with the Mouse GRCm38 reference genome. The feature-ref flag of Cell Ranger was used to generate a gene expression matrix counts alongside the expression of cell surface proteins. The obtained gene-barcode matrix was further analyzed using Seurat v5 (5). Hashtag and CITE-seq antibody counts were separated from the main gene expression (RNA) matrix. RNA data was normalized using the NormalizeData function, the top 2,000 highly variable genes were identified and then scaled using the ScaleData function. Hashtag and CITE-seq data were added as independent assays and normalized using the NormalizeData with CLR transformation. Demultiplexing was performed to identify sample of origin, multiplets exclusion, and cells with undetectable hashtag signal using HTODemux function in Seurat setting "positive quantile = 0.9999". Dead cells and potential outliers were filtered out by excluding cells with more than 5% of mitochondrial genes and 40000 total RNA counts, respectively. Experimental groups were assigned based on the hashtag signals (**Table 2**). Cluster analysis was performed using the FindCluster function in Seurat using 30 principal component and 0.5 resolution. Proportion of clusters across experimental conditions was assessed by using the ClusterDistrBar function of the SeuratExtend package (8). The MAM signature was applied using the AddModuleScore function to the BMDMs dataset. Differentially expressed genes between cluster C6 and the others were performed using the VolcanoPlot function from the SeuratExtend package. Reactome analysis was performed comparing clusters C4 and C6 using the "extracellular matrix organization" gene set. To evaluate the effect of IL-4 priming during efferocytosis, C6 was isolated from the total dataset and only the Apo_cells and IL4+Apo_cells conditions were selected for further analysis. Differentially expressed genes in C6 were identified using the VolcanoPlot function from the Seurat Extend package between Apo_cells and IL4+Apo_cells conditions. Gene set enrichment analysis of the MAM signature was performed using the GSEAplot function of the SeuratExtend package in C6 between Apo_cells and IL4+Apo_cells conditions.

Analysis of human cells after acute myocardial infarction

We analyzed published available data of human heart cells (GSE217494) (9). Only patients with acute myocardial infarction and donor control were processed for downstream analysis. The library of each patient were pre-processed independently for quality control using the same pipeline. Raw count matrices were processed using Seurat. CITE-seq antibody counts were separated from the main gene expression (RNA) matrix. RNA data was normalized using the NormalizeData function, the top 2,000 highly variable genes were identified and then scaled using the ScaleData function. Hashtag and CITE-seq data were added as independent assays and normalized using the NormalizeData function with CLR transformation. Dead cells and potential outliers were filtered out by excluding cells with more than 5% of mitochondrial genes and 30000 total RNA counts, respectively. Data from each patient was independently processed for quality control. Dead cells were filtered out by high expression of mitochondrial genes. After quality control, the libraries from each patient were integrated with harmony and used to assign the experimental groups. Cluster analysis was performed with the FindCluster function from the Seurat package using 25 principal component and 0.6 resolution. Monocytes and macrophages from the total cell pool were extracted for in depth characterization. Clustering analysis was performed using 20 principal components and a 1.0 resolution. Monocytes and macrophages were annotated based on the top30 marker genes calculated with the FindAllMarkers function from the Seurat package. The ClusterDistrBar function from the Seurat extend package was utilized to visualize monocyte and macrophage propotion across conditions. MAM signature was applied to the monocyte and macrophage clusters using the AddModuleScore function in Seurat. We used the human (GSE217494) and the mouse (this study) monocyte and macrophage datasets and performed cross-species analysis using the ClusterFoldSimilarity package (10). The similarities values were exported (**Table 2**) and visualized using the clusterFoldSimilarity() function.

**CurioSeeker Spatial Transcriptomics**

Spatial transcriptomics of an infarcted male C57BL6/J mouse (day 7 post-MI) was performed on a fresh frozen cryosection using the Curio Seeker Spatial Mapping Kit (Curio Biosciences). The heart was snap frozen in liquid nitrogen-cooled isopentane. Using a Cryostat, a 10µm section was placed on the CurioSeeker slide´s capture area and processed for sequencing following the manufacturer´s instruction. Briefly, the tile with the heart tissue was removed and placed in an Eppendorf tube to perform the following step of hybridization to the Seeker tile and reverse transcription. Then, tissue clearing was performed followed by cDNA amplification and library preparation. Sequencing was performed using Novaseq6000 (Illumina). The sequencing data was processed and aligned to GRCm38 using the Curio seeker bioinformatics pipeline (v.1.0.4) to generate a gene count x bead matrix with matched bead barcodes between array and sequenced reads. The bead barcode location files for the tile were provided by Curio biosciences. The gene count x bead barcode matrix and the positional information were loaded into R (v4.2.1) and converted into a Seurat object using the Seurat toolkit (v5.0.1), which was used for downstream analyses. Beads with fewer than 80 detected UMIs were removed. To ensure only beads from actual tissue were retained, we argued that such beads should have a certain amount of neighboring tissue beads within a certain radius. Therefore, neighboring retained beads within a radius of 100 pixels were identified for every bead using the nn2 function of the R package RANN (v2.6.1). Beads with less than 10 neighbouring beads with >80 UMIs were removed. Robust cell type decomposition (RCTD; R package spacexr v2.2.1) (11) was used to deconvolve the bead transcriptomes using a sc/snRNA-seq reference dataset for training, comprising non-myocytes cells from the day 7 time point of the data shown in **Figure 1**, merged with a cluster corresponding to cardiomyocytes obtained from a single-nucleus RNA-seq dataset.

**Statistical analysis**

Statistical analyses were performed using GraphPad Prism version 10. Results are expressed as mean ± s.e.m. For two-group comparisons, normal distribution of the data was assessed by a D'Agostino–Pearson test followed by an unpaired t-test (normally distributed data) or a non-parametric Mann–Whitney test (non-normally distributed data). Data with multiple comparisons were assessed by one-way ANOVA followed by a Holm-Šídák's multiple comparisons test. P values less than 0.05 were considered statistically significant.

### Supplemental Figures

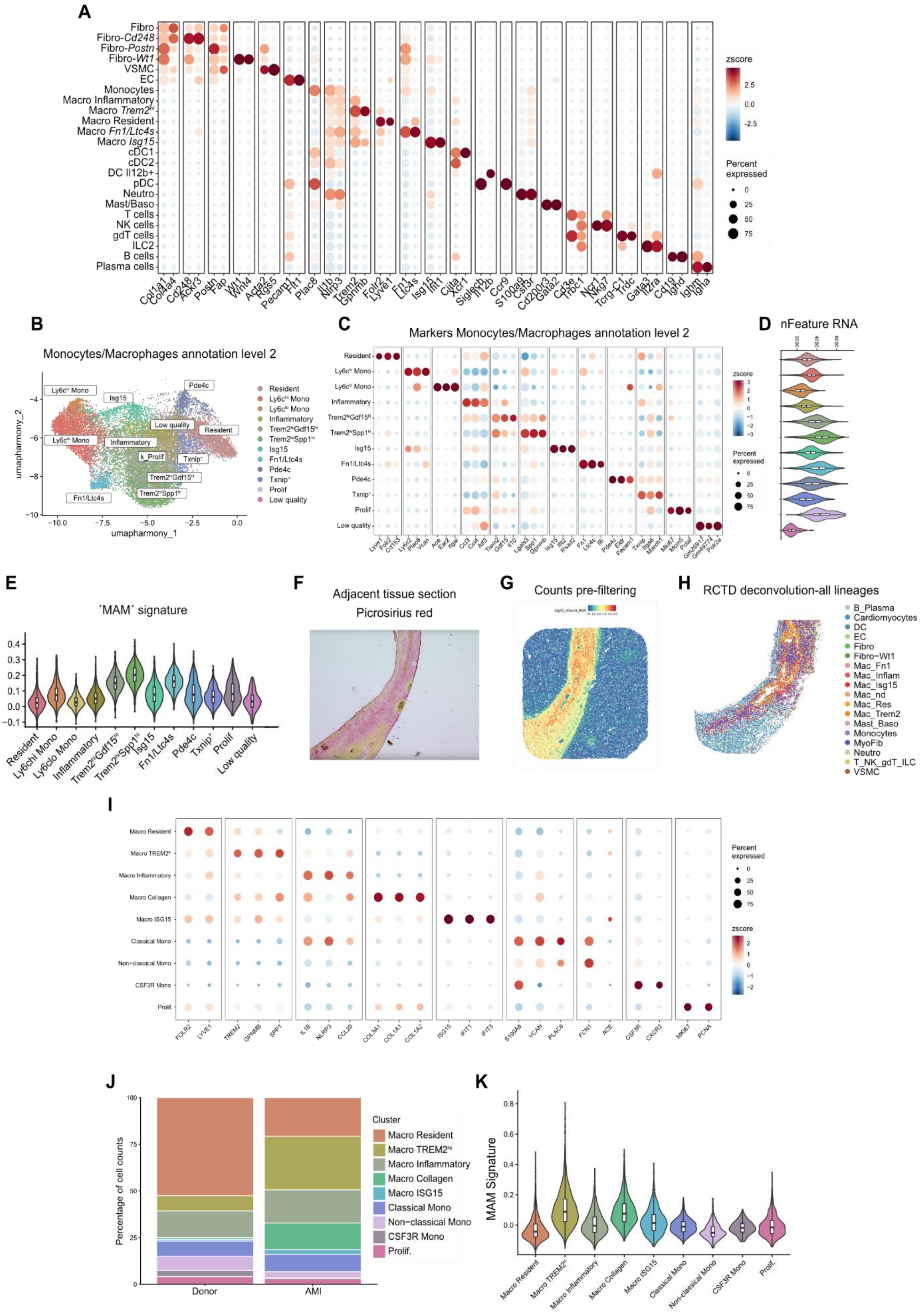

**Supplemental Figure 1**. ***Trem2**$^{hi}$* **macrophages co-localize with myofibroblasts in the infarcted heart**. **A)** Dotplot showing the main marker genes used to identify cell clusters in **Figure 1b**. **B)** UMAP plot of monocytes and macrophages extracted from **Figure 1b** and reclustered. **C)** Dotplot showing the main markers used to annotate monocytes and macrophages. **D)** Expression of RNA features in monocytes and macrophages subclusters. **E)** Violin plot showing the MAM signature score in macrophage subsets from panel **B**. **F)** Representative picrosirius red staining picture of serial heart section at day 7 post-MI used for the Curio Seeker spatial transcriptomics experiment. **G)** RNA counts projected onto the Curio Seeker spatial transcriptomics data before pre-filtering. **H)** Curio seeker data after filtering of low RNA spots showing the localization of cell type identified with RCTD deconvolution. **I)** Dotplot with the main markers used to identify human cardiac monocyte and macrophage populations of **Figure 1j**. **J)** Monocyte and macrophage clusters distribution. **K)** Expression of the MAM signature score in human cardiac monocyte and macrophage clusters from **Figure 1j**.

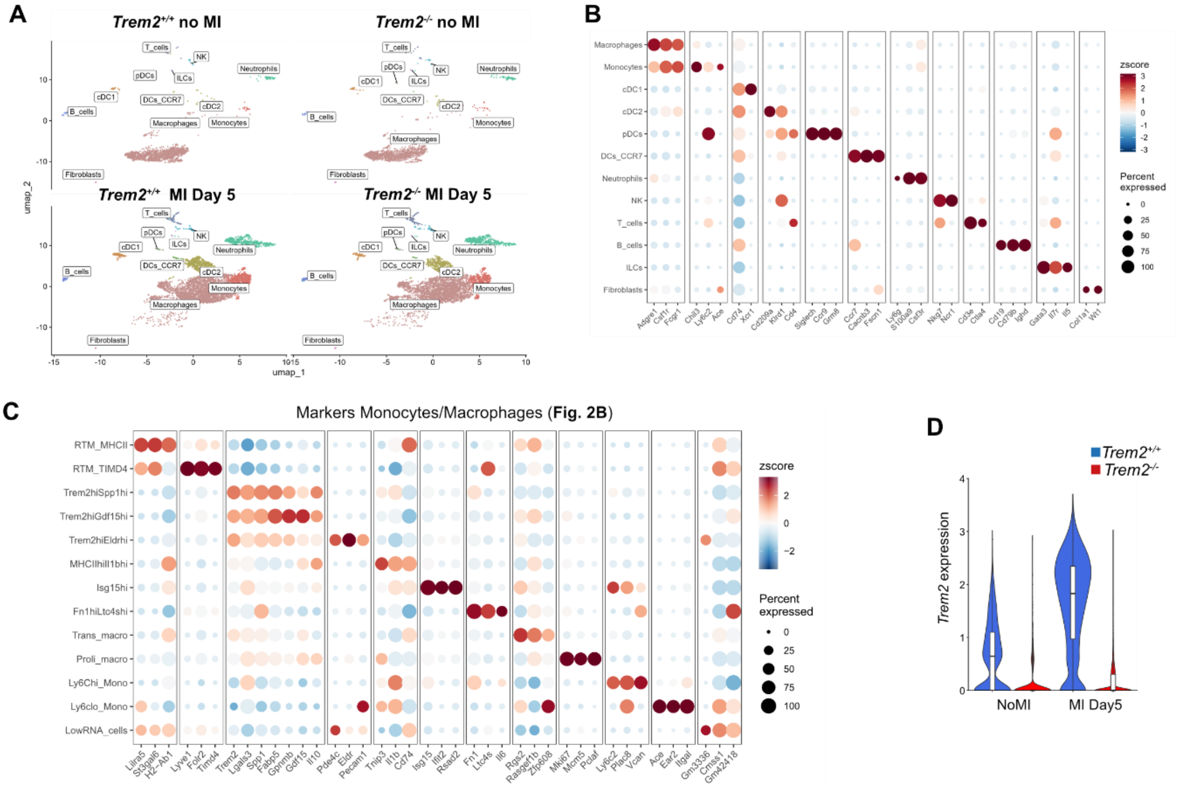


**Supplemental Figure 2**. ***Trem2* drives accumulation of monocyte-derived macrophages with a pro-scarring matrisome-associated gene expression profile in the infarcted heart**. **A)** UMAP plot of cardiac leukocytes from *Trem2*$^{+/+}$ and *Trem2*$^{-/-}$ mice at day 5 post-MI. *Trem2*$^{+/+}$ No MI n= 2, *Trem2*$^{-/-}$ No MI n = 3, *Trem2*$^{+/+}$ MI Day 5 n= 6, *Trem2*$^{-/-}$ MI Day 5 n = 6; **B)** Dotplot showing the main marker genes used to identify the cell identity in **a**; **C)** Dotplot showing the main marker to identify monocyte and macrophage subclusters ; **D)** *Trem2* expression in *Trem2*$^{+/+}$ and *Trem2*$^{-/-}$ mice at baseline and days 5 post-MI. *Trem2*$^{+/+}$ MI Day 5 n= 6, *Trem2*$^{-/-}$ MI Day 5 n = 6.

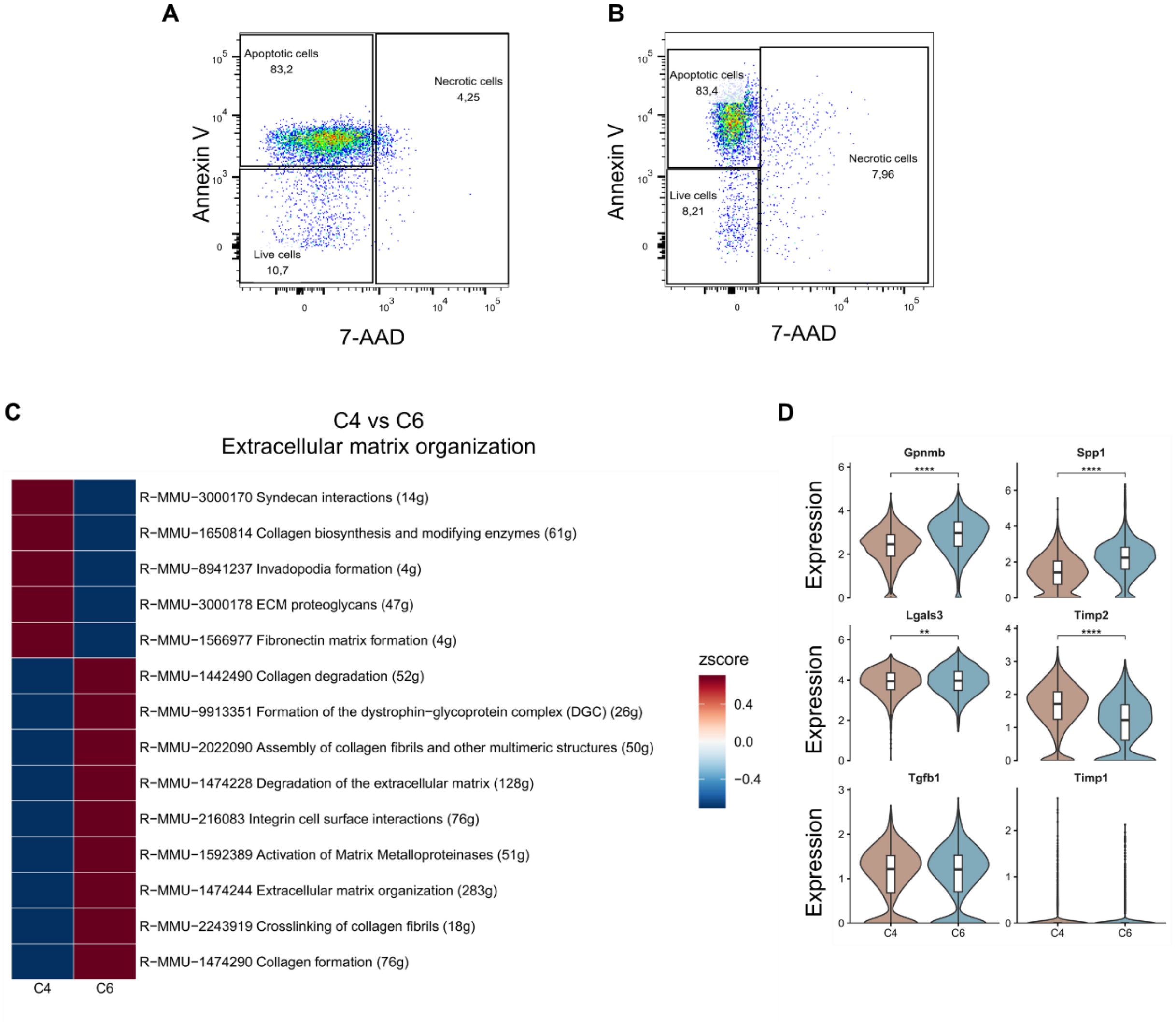


**Supplemental Figure 3**. **Efferocytosis drives acquisition of the pro-scarring macrophage signature A)** Representative flow cytometry plot of Annexin V and 7AAD staining in apoptotic neutrophils; **B)** Representative flow cytometry plot of Annexin V and 7AAD staining in apoptotic Jurkat cells; **C)** Reactome analysis of "extracellular matrix organization" pathways in C4 and C6; **D)** Violinplot of the indicated genes in C4 and C6.
